# β-Catenin tumour-suppressor activity depends on its ability to promote Pro-N-Cadherin maturation

**DOI:** 10.1101/571810

**Authors:** Antonio Herrera, Anghara Menendez, Blanca Torroba, Sebastian Pons

## Abstract

Neural stem cells (NSCs) form a pseudostratified, single-cell layered epithelium with a marked apico-basal polarity. In these cells, β-Catenin associates with classic cadherins in order to form the apical adherens junctions (AJs). We previously reported that oncogenic forms of β-Catenin (sβ-Catenin) maintain neural precursors as progenitors, while also enhancing their polarization and adhesiveness, thereby limiting their malignant potential. Here we show that β-Catenin can bind to phosphorylated Pro-N-Cadherin, promoting the excision of the propeptide and its maturation into N-Cadherin in the trans-Golgi network (TGN). Moreover, β-Catenin-assisted maturation of Pro-N-Cadherin is required for the formation of the AJs and for them to recruit other apical complex (AC) components like aPKC, and accordingly, to establish apico-basal polarity. Notably, we show that NSCs expressing unprocessed Pro-N-Cadherin invade the ventricle and they breach the basement membrane to invade the surrounding mesenchyme. Hence, we propose that the tumour-suppressor activity of sβ-Catenin depends on it promoting Pro-N-Cadherin processing.

## INTRODUCTION

Neural stem cells (NSCs) form a pseudostratified, single-cell layered epithelium, extending from the ventricle to the basal lamina and displaying marked apico-basal polarity. The proteins present at the apical pole of NSCs are collectively called the apical complex (AC) and they were initially described in the cerebral cortex, forming three concentric sub-domains with discrete functions: the fate-determining factors (PAR3, aPKC, Prominin1) confined within or close to the apical membrane; the zonular proteins (ZO1, Afadin and Actin) occupying an intermediary position; and the junctional complexes (N-Cadherin, α-Catenin and β-Catenin) located in the sub-apical domain (Marthiens & ffrench-Constant, 2009). However, although different AC proteins are enriched in these domains, there is extensive overlap in their distributions. During development, NSCs can initially be identified by the expression of Sox2 or CD133/prominin1 (Zhu et al, 2009), and these cells proliferate symmetrically in a self-expanding mode. Later on, and in association with the onset of neurogenesis, their mode of division changes to generate the first committed neurons (Gotz & Huttner, 2005; Saade et al, 2013). In these neurogenic divisions, post-mitotic NSCs dampen their expression of N-Cadherin and they detach from the proliferative ventricle (Das & Storey, 2014; Rousso et al, 2012). Thus, it is critical to coordinate NSC differentiation and delamination in order to maintain the architecture of the nervous system and to avoid the initiation of certain pathologies.

β-Catenin mediates canonical WNT signalling, stimulating Tcf dependent transcription (Grigoryan et al, 2008; Nelson & Nusse, 2004). However, β-Catenin also plays important roles in epithelial cell polarity, for example associating with classic cadherins through its armadillo domains and thereby contributing to Adherens Junction (AJ) formation (Baum & Georgiou, 2011). Classical cadherins share a basic structure and role in adherence, and they are named on the basis of the tissue in which they are mainly expressed (i.e.: E-Cadherin in epithelial tissue or N-Cadherin in neural tissue). These glycoproteins are synthesized as precursors that must undergo a series of post-translational modifications to become functional at AJs. The association between the mature form of classic cadherins and β-Catenin has been widely studied, however, the relationship between β-Catenin and the precursor Pro-Cadherins is still poorly understood.

Although apico-basal polarity is normally lost during the acquisition of a malignant phenotype (Al-Masri & McCaffrey, 2015), there is little evidence as to how polarity proteins regulate malignancy. It has been reported that a loss of Par3 promotes tumorigenesis and metastasis in breast cancer (McCaffrey et al, 2012; Xue et al, 2013), and that most aPKC mutations in cancer are loss-of-function mutations (Antal et al, 2015). This is consistent with the fact that genes that induce the epithelial to mesenchymal transition (EMT) suppress E-Cadherin expression as part of their genetic program (Thiery et al, 2009). Likewise, the invasive behaviour of chemo-resistant cells derived from breast tumours, has recently been associated with the expression of non-adherent forms of Pro-N-Cadherin at their cell surface (Nelson et al, 2016). Notably, one of the mechanisms by which E-cadherin prevents cell growth and transformation is by recruiting β-Catenin to AJs, blocking its transcriptional activity and enhancing cell-cell cohesion (Gottardi et al, 2001; Stockinger et al, 2001). Thus, cell polarity and AJs must themselves be considered as tumour suppressors.

Activation of the WNT signalling, mostly due to weaker β-Catenin degradation (Reya & Clevers, 2005), has been observed in many brain cancers, including medulloblastoma, the most common paediatric malignant primary brain tumour. Consequently, the transcriptional and structural activities of β-Catenin will be enhanced in these tumours. Medulloblastomas are classified into four subtypes on the basis of their molecular signature: SHH, WNT, Group 3 and 4 (Taylor et al, 2012). In the WNT subtype of medulloblastoma stable forms of β-Catenin (sβ-Catenin) and strong *MYC* expression are evident, and because it rarely metastasizes it is the least aggressive of the four subtypes of medulloblastomas (Taylor et al, 2012). By contrast, Group 3 medulloblastomas overexpress or carry *MYC* amplifications and they fail to activate the WNT pathway. These medulloblastomas have the worst prognosis due to their very high incidence of spinal metastasis (Pei et al, 2012; Taylor et al, 2012) and in these metastases, EMT genes are expressed more strongly than in the primary tumour (Kahn et al, 2018). Notably, β-Catenin overexpression is associated with a favourable outcome in these tumours, even when areas of large-cell anaplastic (LCA) histology are evident, a poor prognostic marker (Ellison et al, 2011). Moreover, sβ-Catenin expression significantly augments the survival of cells derived from the SHH medulloblastoma mouse model (Poschl et al, 2014). Similarly, a mouse model of WNT-type medulloblastoma develops aberrant groups of cells in the dorsal brainstem during the foetal period, these rarely progressing to invasive tumours unless other genes like *TP53* or *PIK3CA* are also mutated (Gibson et al, 2010; Robinson et al, 2012). In summary, there is significant evidence of the tumour-suppressor role of β-Catenin in medulloblastoma.

We previously reported that the activity of sβ-Catenin in the developing neural tube maintained cells as progenitors, although it also increased polarization and adhesiveness, limited proliferation (Herrera et al, 2014), and reduced delamination and migration (Rabadan et al, 2016). Thus, we hypothesized that by inducing the expression of genes like *MYC*, which contributes to the maintenance of stemness, sβ-Catenin acts as an oncogene as well as a tumour suppressor, promoting the formation of enlarged ACs that limit the invasive properties of sβ-Catenin expressing cells. Here, a more in depth analysis of the molecular mechanisms by which sβ-Catenin promotes AC formation in NSCs has been performed. We show that sβ-Catenin binds to phosphorylated Pro-N-Cadherin, promoting the excision of the propeptide and its final maturation into N-Cadherin in the trans-Golgi network (TGN). We also show that β-Catenin-aided maturation of Pro-N-Cadherin is an absolute requirement for the formation of AJs and for the recruitment of other AC components like aPKC, and accordingly, for the establishment and maintenance of apico-basal polarity. The effect of sβ-Catenin on Pro-N-Cadherin maturation is independent of transcription and remarkably, we demonstrate that the persistence of Pro-N-Cadherin leads to the loss of the apical AJs in proliferating NSCs. As a result, these cells either invade the ventricle or prematurely delaminate due to the retraction of the apical process.

Finally, we demonstrate that faulty delamination of NSCs leads to their accumulation in the mantle zone, where they provoke neural tube deformities and finally, breach the basal membrane to invade the surrounding mesenchyme.

## RESULTS

### β-Catenin induces the apical localization of aPKC through its binding to N-Cadherin

We previously demonstrated that oncogenic forms of β-catenin (sβ-Catenin) induce aberrant growth of the neuroepithelium by enhancing the accumulation of AC proteins at the apical pole of NSCs (Herrera et al, 2014). This phenotype required aPKCι and it could not be reproduced by constitutive activation of the Wnt pathway through VP16-TCF3. As a result, we hypothesized that the structural and transcriptional activities of β-catenin were required to deliver N-Cadherin and aPKC to the AC. To decipher the contribution of each of these activities to the formation/maintenance of the AC, and consequently to the integrity of the neural epithelium, we studied the distribution of N-Cadherin and aPKC under different conditions where the structural and transcriptional activities of β-Catenin were modified independently (Fig. 1A-C, and Fig. EV1). From images of aPKC and N-Cadherin expression at 48 hpe (hours post-electroporation: Fig. 1A), the apico-basal distribution of these proteins was studied. All images were normalized to 100 pixels (the first fifteen considered apical and the remaining 85 basolateral), and the pixel intensity profiles for aPKC and N-Cadherin were plotted, calculating the area under the curve for the apical (∫_A_) and basolateral (∫_BL_) regions (Fig. 1B). Finally, and as an indicator of neuroepithelial polarization, we calculated the apical/basolateral ratios (A/BL) of aPKC and N-Cadherin, normalized to their respective controls in order to facilitate a comparison between these two proteins: (∫_A_/∫_BL_)^Control^/(∫_A_/∫_BL_)^Treatment^ (Fig. 1C). In addition, we calculated the incidence of invaginations and lumen invasion associated with the different treatments (Fig. 1D).

**Figure 1.**
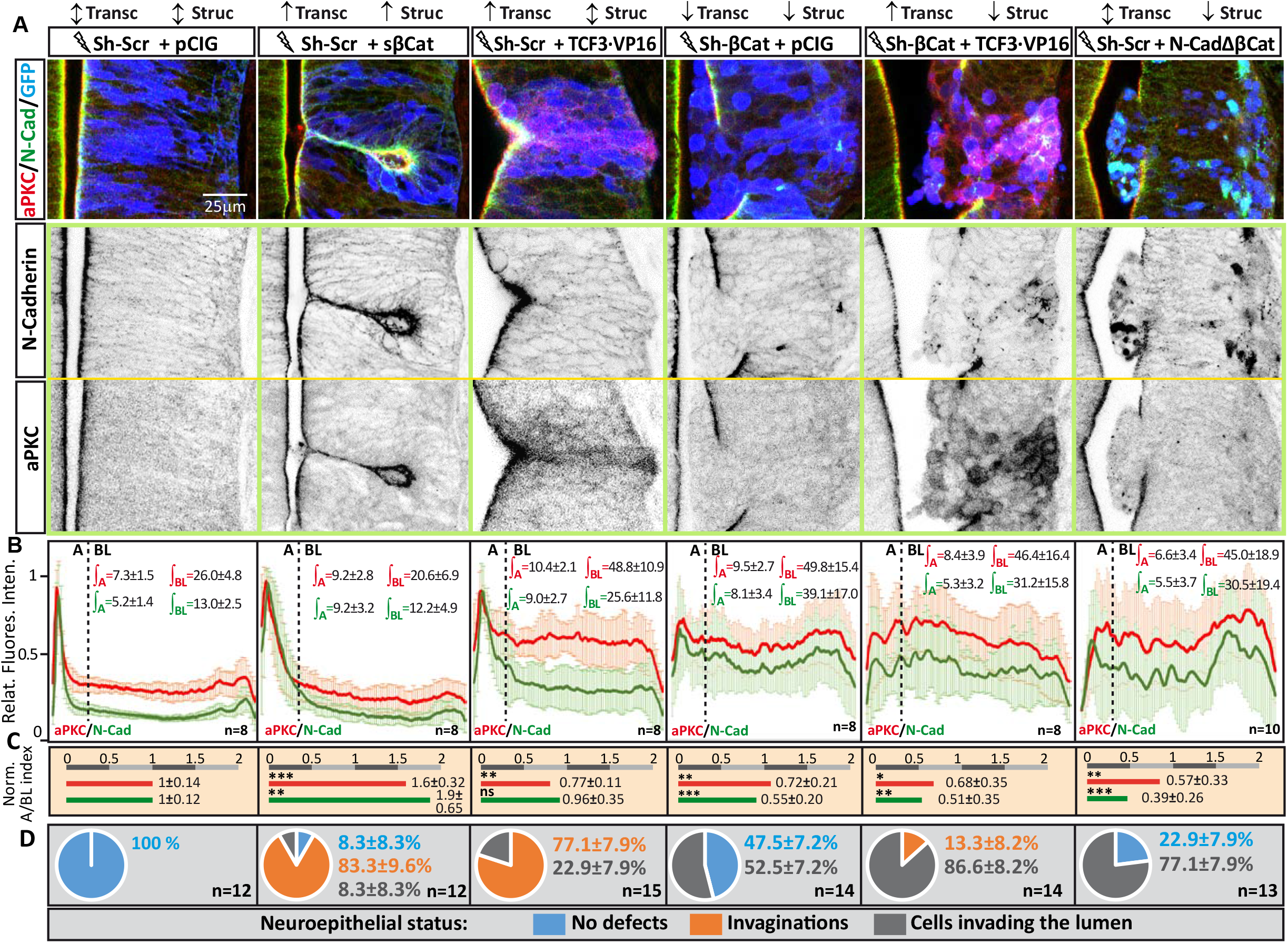
β-Catenin induces apical localization of aPKC through its binding to N-Cadherin. **(A)** HH12 Chicken neural tubes were transfected with different molecular tools intended to independently manipulate the transcriptional (Transc) and structural (Struc) activities of β-Catenin. The figure shows representative images of the transfected neural tubes used for quantification. The effect caused by each combination at 48 hpe is indicated: ţincreased activity, ↓decreased activity, ↕ no change. pCIG indicates the empty vector, and Sh-Scr and Sh-βCat the scrambled and the β-Catenin short-hairpin inhibitory RNAs, respectively. TCF3·VP16 is a constitutive activator of TCF dependent transcription and N-CadherinΔβCat is a N-Cadherin mutant lacking the β-Catenin binding domain. **(B)** The apico-basal distance in the images with a similar level of transfection was normalized to 100 pixels, with the first 15 pixels arbitrarily considered to be apical and the remaining 85 basolateral. The line-plots represent the pixel intensity profiles for aPKC and N-Cadherin, and ∫_A_ and ∫_BL_ represent the values of the area-under-the-curve for the apical and basolateral regions, respectively. **(C)** The bar charts shown represent the apical/basolateral ratios (A/BL) of aPKC and N-Cadherin expression, normalized to their respective controls ((∫_A_/∫_BL_)^Control^/(∫_A_/∫_BL_)^Treatment^). **(D)** The pie charts represent the percentage of growth aberrations observed in each condition (mean ± SEM): *ns*, non-significant; \**p* <0.05; \*\**p* <0.01; \*\*\**p* <0.001. Lightning bolts denote transfection.

The presence and expression of sβ-Catenin (“structural and transcriptional up”) induced the formation of larger ACs that contained more aPKC and N-Cadherin, causing a characteristic tissue invagination (Herrera et al, 2014). By contrast, β-Catenin silencing with Sh-β-Catenin (“structural and transcriptional down”) enhanced the amount of N-Cadherin at the lateral membrane, favouring a round shape of NSCs and inducing the invasion of the ventricle by groups of transfected cells. Hence, the deficit in β-Catenin impairs AJ function, compromising the cohesion of the epithelium in a manner consistent with the phenotype reported for the β-Catenin KO (Valenta et al, 2011).

Transfection of VP16-TCF3 (“transcriptional up”) induced aPKC and N-Cadherin accumulation at the AC, provoking small invaginations similar to those produced by sβ-Catenin transfection. However, although VP16-TCF3 stimulated the expression of aPKC similarly to sβ-Catenin, the A/BL ratio of aPKC was reduced because it mostly remained in the basolateral compartment. Interestingly, co-expression of VP16-TCF3 with Sh-β-catenin (“structural down” and “transcriptional up”) disrupted the epithelium and the formation of ACs of NSCs, generating an unpolarised cell mass in the lumen. This response highlights the importance of β-Catenin’s structural activity for aPKC accumulation in the AC. Notably, a very similar phenotype was obtained with a mutant N-Cadherin that lacks the β-Catenin binding domain (N-CadherinΔβCat). In brief, β-Catenin transcriptional activity induces the expression of proteins that like aPKC, are relevant for AC formation and cell polarization. However, the structural function of β-Catenin is an absolute requirement to construct ACs, to maintain apico-basal polarity and to ensure the integrity of the neuroepithelium. Moreover, the structural activity of β-Catenin involves its binding to N-Cadherin.

### β-Catenin binds to N-Cadherin in the Golgi apparatus of NSCs to form sub-apical Adherens Junctions (AJs)

To better understand how the structural activity of β-Catenin participates in the maintenance of apico-basal polarity, we studied the interactions among N-Cadherin, aPKCι and β-Catenin. We individually electroporated ST-tagged forms of these three molecules into HH-12 chicken embryos and we then purified them at 24 hpe on Streptactin affinity columns, studying the associations of their endogenous partners in western blots (Fig. 2A,B). N-Cadherin-ST and sβ-Catenin-ST efficiently co-purified the other two proteins (Fig. 2C,D), while aPKCι-ST preferably co-purified with β-Catenin rather than N-Cadherin (Fig. 2E). Notably, neither β-Catenin nor aPKCι co-purified with N-CadherinΔβCatBD-ST, a mutant N-Cadherin lacking the β-Catenin binding domain, indicating that aPKCι interacts with N-Cadherin through β-Catenin (Fig. 2F). As already reported in the telencephalon (Marthiens & ffrench-Constant, 2009), the distribution of aPKC was more apical than that of N-Cadherin/β-Catenin in the AC, although a significant amount of aPKC co-localized with N-Cadherin/β-Catenin at the boundary between the apical and sub-apical regions (Fig. 3A,B,A’,B’). Similarly, although some actin expression was detected apical to the N-Cadherin domain, most of the actin in the AC was detected within the domain of N-Cadherin expression (Fig. EV2).

**Figure 2.**
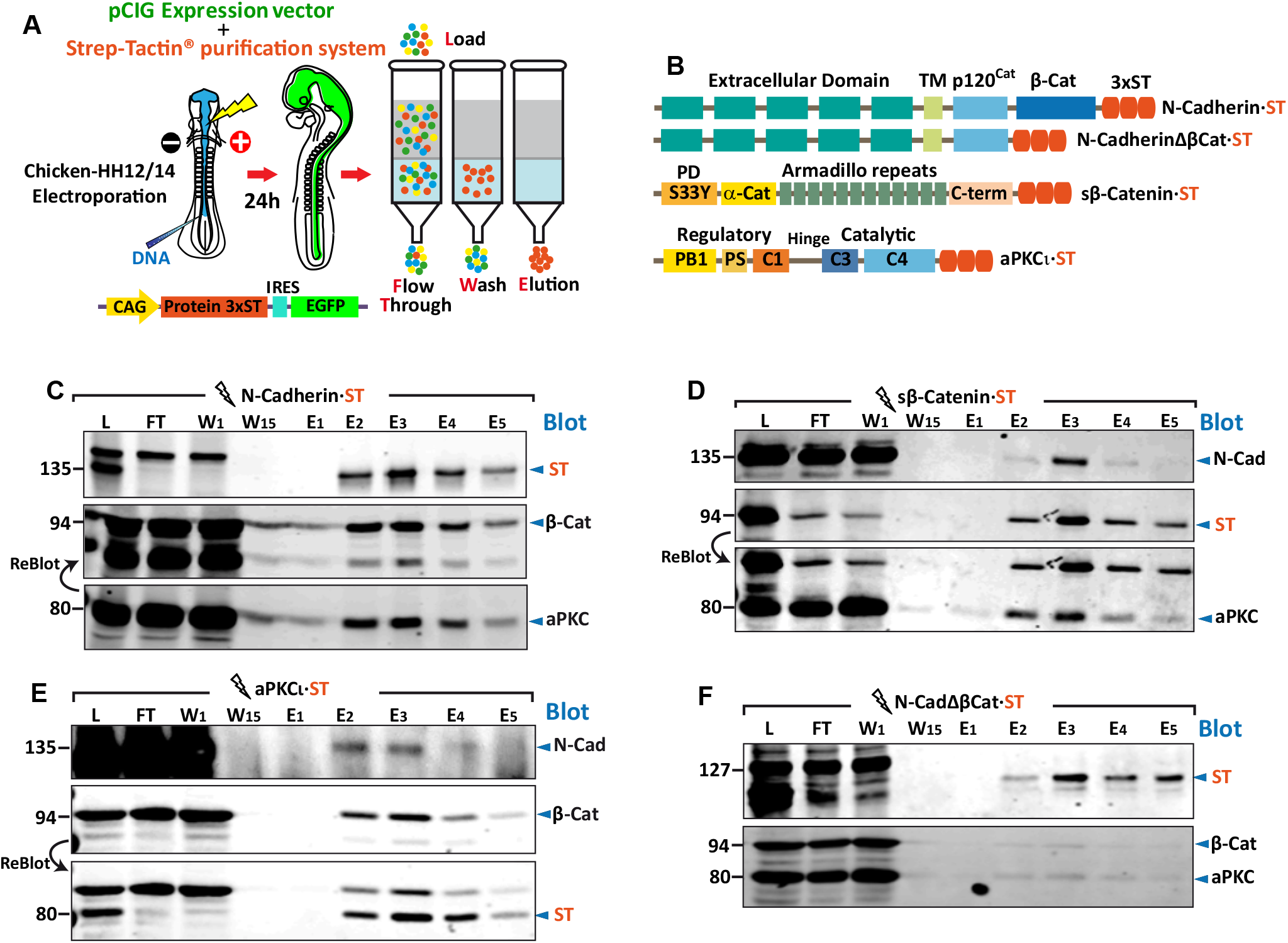
N-Cadherin forms complexes with β-Catenin and aPKC. **(A)** Representation of the *in ovo* electroporation and Streptactin-Streptag^®^ purification procedures, and **(B)**, scheme of the four ST-tagged molecules used. **(C-F)** The different ST-tagged proteins were electroporated into HH12 chicken neural tubes, and the ST-tagged and associated endogenous proteins were purified on Streptactin columns at 24 hpe: L, Lysate; FT, Flow Through; W, Wash; E, Elution. **(C)** Electroporation and purification of N-Cadherin·ST probed with antibodies against ST (N-Cadherin), aPKCι and β-Catenin, note that the β-Catenin image is a re-probing of the aPKCι blot. **(D)** Electroporation and purification of sβ-Catenin·ST probed with antibodies against N-Cadherin, ST (sβ-Catenin) and aPKCι, the aPKCι image is a re-probing of the ST blot. **(E)** Electroporation and purification of aPKCι·ST probed with antibodies against N-Cadherin, β-Catenin and ST (aPKCι), the ST blot is a re-probing of the β-Catenin blot. **(F)** Electroporation and purification of N-CadherinΔβCatST probed simultaneously with antibodies against ST (N-CadherinΔβCat) and β-Catenin or anti PKCι. Note that neither β-Catenin nor PKCι co-purify with N-CadherinΔβCatST. Lightning bolts denote transfection.

To assess the degree of co-localization, we calculated the Manders’ correlation coefficients for aPKC/N-Cadherin and β-Catenin/N-Cadherin. The coefficient for aPKC/N-Cadherin was significantly higher in the AC than in the rest of the cell (Basolateral), where it was close to zero (0.41±0.04 versus 0.03±0.01). Hence, aPKC appears to co-localize with N-Cadherin almost exclusively in the AC (Fig. 3C,D). By contrast, while the β-Catenin/N-Cadherin coefficient was significantly higher in the AC than in the whole cell (0.90±0.02 versus 0.63±0.02), in this case the co-localization was also most evident in the region between the nucleus and the AC. In this space, the staining of N-Cadherin and β-Catenin was punctuate and the degree of co-localization seemed to increase in the proximity of the AC. In the developing brain cortex, the radial glia have a very elongated Golgi apparatus located apical to the nucleus and vesicular trafficking takes place perpendicular to the apico-basal axis (Taverna et al, 2016). We observed a similar Golgi distribution in the NSCs of the developing neural tube (Fig. 3E), and calculated the co-localization index of N-Cadherin/β-Catenin in the Golgi and in the AC-to-Golgi regions (Fig. 3E’). Notably, the co-localization index was significantly higher at the AC-to-Golgi than in the Golgi apparatus itself (0.46±0.03 versus 0.24±0.03: Fig. 3F,G). These data strongly suggests that β-Catenin associates with N-Cadherin in the Golgi during its journey to the AJs, while aPKC interacts with the complex exclusively in the AC.

**Figure 3.**
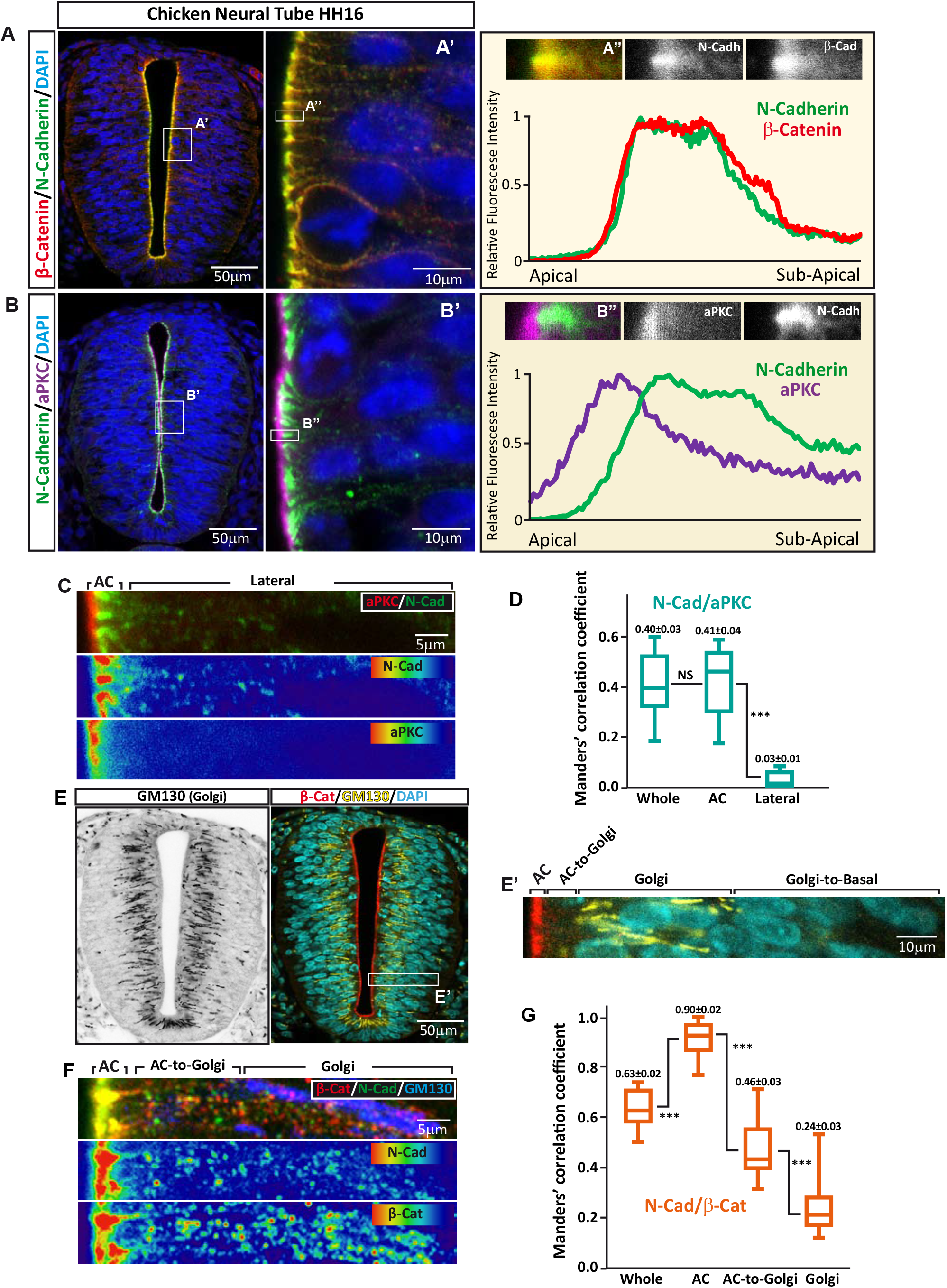
N-Cadherin co-localizes with β-Catenin in the Golgi-to-AC space, and with β-Catenin and aPKC in the AC. **(A)** HH16 chicken neural tubes were stained with antibodies against β-Catenin (Red), N-Cadherin (Green) and DAPI (blue), the area labelled as A’ is enlarged in the right-hand panel. The graph shows the pixel intensity profile for N-Cadherin and β-Catenin in the area labelled as A”;, with the area quantified shown above. The two channels are displayed separately in grey scale for clarity. **(B)** As in A but using antibodies against N-Cadherin and aPKCι. **(C)** High magnification image of the apico-lateral region of HH16 neural tubes stained with aPKCι (Red) and N-Cadherin (Green), the two lower panels show the N-Cadherin and aPKCι channels separately using a High-Low false colour index. **(D)** Box-plot showing the Manders’ correlation coefficients for N-Cadherin and aPKCι. **(E)** Grey-scale and three channel image of the Golgi apparatus (stained with GM130) in the HH16 neural tube, β-Catenin and DAPI are used as landmarks. The area labelled as E’ is enlarged on the right and is used to define 4 sub-cellular regions: AC, AC-to-Golgi, Golgi, and Golgi-to-basal. **(F)** High magnification image of the apico-lateral region of HH16 neural tubes stained for β-Catenin (Red), N-Cadherin (Green) and GM10 (Blue). The two lower panels show the N-Cadherin and β-Catenin channels separately using a High-Low false colour index. **(G)** Box-plot showing the Manders’ correlation coefficients for N-Cadherin and β-Catenin calculated for the areas described in fig E’: NS, not significant; \**p* <0.05; \*\**p* <0.01; \*\*\**p* <0.001.

### β-Catenin drives the apical localization of N-Cadherin

N-cadherin accumulates at the apical AJs of NSCs, whereas actin accumulates at both the apical and basal poles (Fig. 4A,B). While N-Cadherin must interact with β-Catenin to accumulate in the ACs (Fig. 1A-C), it was not clear whether β-Catenin promoted the stability/recycling of the pre-existing N-Cadherin, or if it facilitated the transport/maturation of the newly synthetized protein. We also found that an important proportion of the N-Cadherin-GFP fusion protein remained in the basolateral compartment (Fig. 4C,D), which we believe might reflect a more extreme molar ratio between the endogenous β-Catenin and the exogenous N-Cadherin-GFP. Indeed, sβ-Catenin expression induced the translocation of N-Cadherin-GFP from the basolateral compartment to the AC (Fig. 4C,D). This “apicalizing” effect of sβ-Catenin was also evident for actin (Fig. 4C,E), possibly because sβ-Catenin induced an enlargement of the ACs that consequently recruited more apical actin filaments (Herrera et al, 2014). These results suggested a role for β-Catenin in the transport or/and maturation of N-Cadherin.

**Figure 4.**
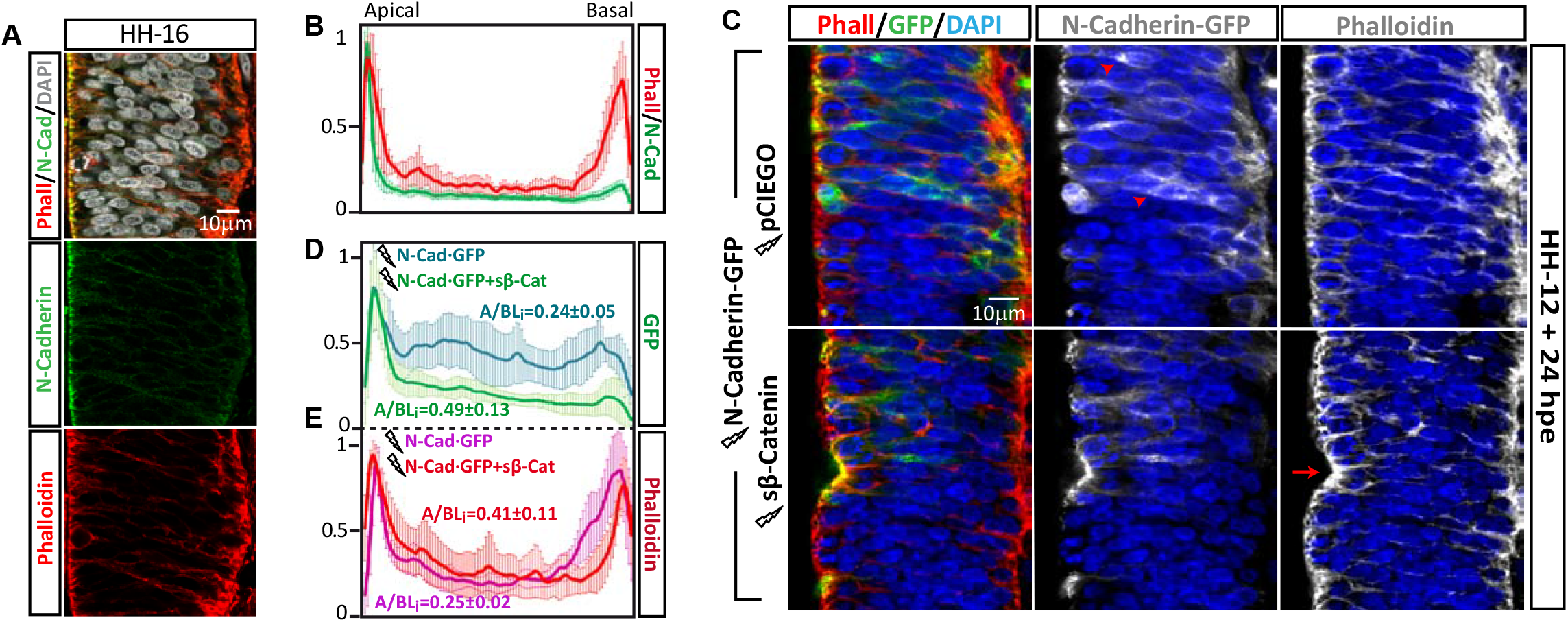
sβ-Catenin promotes the accumulation of GFP-N-Cadherin in the AC. **(A,B)** Representative images and pixel intensity profiles of HH16 chicken neural tube slices stained with Phalloidin (detects f-Actin, Red), anti-N-Cadherin (Green) and DAPI (Grey). **(C-E)** Representative images and pixel intensity profiles of HH12 chicken neural tubes at 24 after electroporation with N-Cadherin·GFP, with or without sβ-Catenin. D and E compare the N-Cadherin·GFP and Actin distributions, respectively. Lightning bolts denote transfection.

To further assess this hypothesis we used HEK-293 cells, an epithelial kidney cell line that does not express endogenous N-Cadherin. In these cells, most of the transfected chicken N-Cadherin·ST accumulated close to the nucleus in what were presumably secretory vesicles (Fig. 5A). Notably, transfected N-Cadherin·ST relocated to the intercellular junctions when sβ-Catenin was expressed in these cells (Fig. 5A), whereas sβ-Catenin failed to redistribute the mutant N-Cadherin lacking the β-Catenin binding domain (Fig. 5B). Indeed, sub-cellular fractionation of HEK-293 cells demonstrated that sβ-Catenin enhanced the proportion of N-Cadherin in the plasma membrane fraction relative to the internal membranes (Fig. 5C,D). By contrast, N-CadherinΔβCat·ST was mainly found in the internal membrane fraction and it did not relocate to the plasma membrane following sβ-Catenin expression (Fig. 5C,D). In short, sβ-Catenin favours the accumulation of N-Cadherin at the plasma membrane, whereas insufficient or defective β-Catenin binding appears to induce the stacking of N-Cadherin at internal membranes.

**Figure 5.**
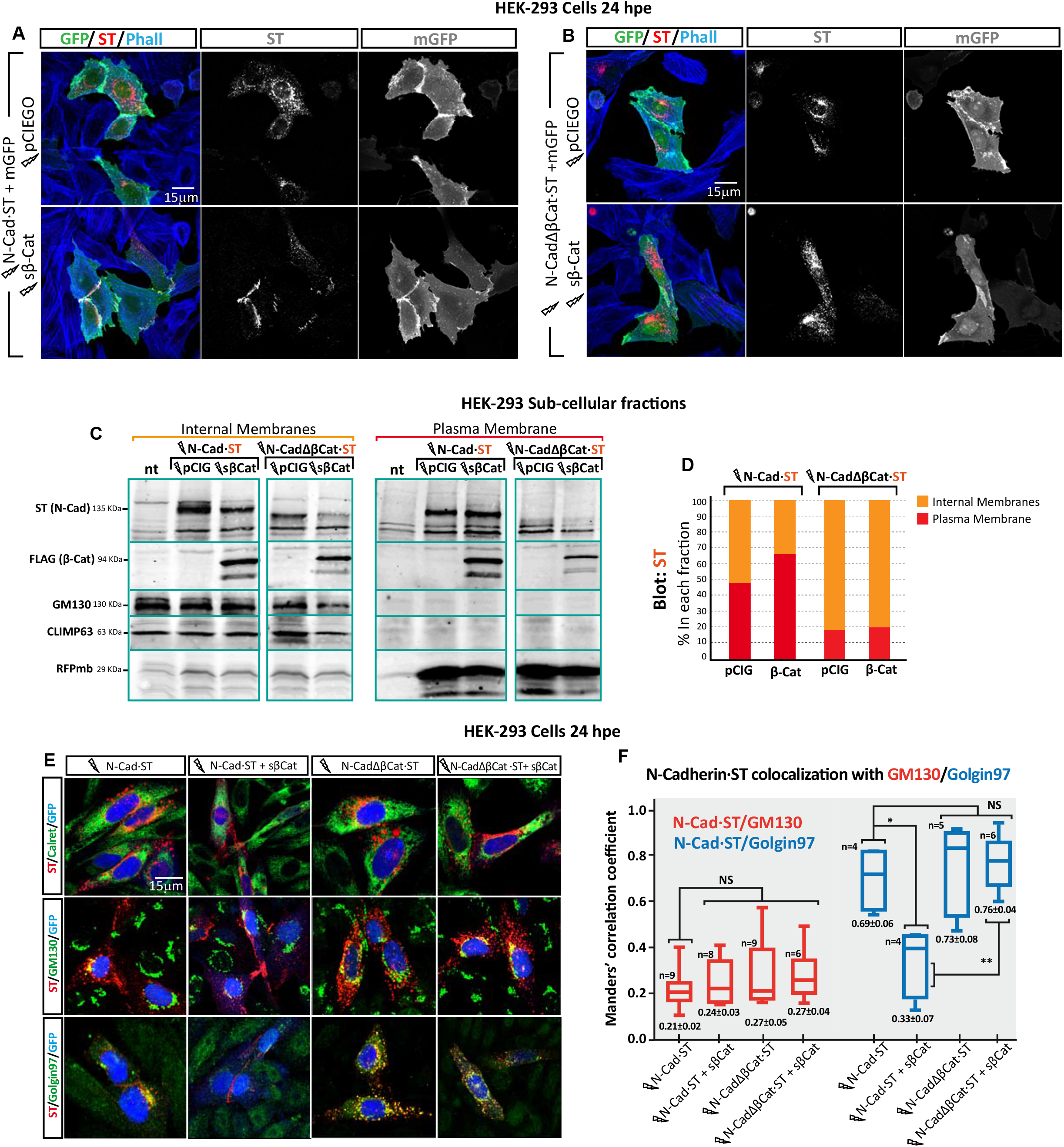
β-Catenin promotes the delivery of N-Cadherin to the intercellular junctions. **(A)** HEK-293 cells 24 hpe with N-Cadherin, with or without sβ-Catenin. Membrane targeted GFP was used to define the plasma membrane of transfected cells and the cells were stained with anti-N-Cadherin (Red) and Phalloidin (f-Actin, Blue). **(B)** A similar experiment as in A but with N-CadherinΔβCat. **(C)** Fractions enriched in internal membranes or plasma membranes were generated from HEK-293 cells transfected with N-Cadherin or N-CadherinΔβCat plus sβ-Catenin or the control vector. The relative amount of N-Cadherin and N-CadherinΔβCat in each fraction was studied in western blots. Climp 63 and GM130 were used as endoplasmic reticulum (ER) and Golgi markers, respectively, and transfected membrane-targeted RFP was used as a plasma membrane marker. **(D)** Quantification of the relative amounts of N-Cadherin and N-CadherinΔβCat in the two fractions (representative of three independent blots). **(E)** HEK-293 cells transfected with N-Cadherin or N-CadherinΔβCat plus the sβ-Catenin or control vector, stained with anti N-Cadherin (Red) and an ER (Calreticulin), Golgi (GM130) or Trans-Golgi/TGN (Golgin 97) marker in green. Nuclear GFP expression indicative of transfection is shown in Blue. **(F)** Box-plot showing the Manders’ correlation coefficients for N-Cadherin and GM130 (Red boxes) or Golgin97 (blue boxes), calculated from cultures in E: NS, not significant; \**p* <0.05; \*\**p* <0.01; \*\*\**p* <0.001. Lightning bolts denote transfection.

### β-Catenin deficiency induces N-Cadherin stacking in the TGN

As indicated above, when chicken N-Cadherin·ST was expressed in HEK-293 cells it was mainly recovered from the internal membrane fraction unless sβ-Catenin was co-expressed. To identify the step at which N-Cadherin trafficking was blocked, we transfected N-Cadherin·ST or N-CadherinΔβCat·ST in the presence or absence of sβ-Catenin, double staining the cells for ST and Calreticulin (endoplasmic reticulum), GM130 (Cys-Golgi) or Golgin97 (TGN: Fig. 5E). Neither N-Cadherin·ST nor N-CadherinΔβCat·ST co-localized with Calreticulin, in the presence or absence of sβ-Catenin (Manders’ coefficient not shown). Although both N-Cadherin·ST and N-CadherinΔβCat· ST co-localized with GM130, this co-localization was weak in both cases (Manders’ coefficients ranging from 0.21±0.02 to 0.27±0.05), and no significant differences were observed between the control and sβ-Catenin transfected cells (Fig. 5F). By contrast, both N-Cadherin·ST and N-CadherinΔβCat·ST co-localized strongly with Golgin97 (Manders’ 0.69±0.06 and 0.73±0.08). Interestingly, the co-localization of N-Cadherin·ST and Golgin97 was significantly dampened after sβ-Catenin transfection (Manders’ 0.69±0.06 to 0.33±0.07), while this did not affect the co-localization of N-CadherinΔβCat·ST with Golgin97 (Manders’ 0.69±0.06 for control and 0.76±0.04 for β-Catenin). It is worth noting that N-Cadherin accumulation in the TGN was accompanied by a rise in the expression of Golgin97, consistent with protein accumulation in the TGN (Fig. 5E). In brief, a lack of β-Catenin caused N-Cadherin to accumulate in trans Golgi vesicles, a situation that was reverted by the expression of sβ-Catenin which facilitated the delivery of N-Cadherin to intercellular junctions.

### Pro-N-Cadherin must interact with β-Catenin to excise the propeptide but not for its glycosylation

Like most membrane proteins, the Pro-N-Cadherin amino acid sequence is assembled in the ER and it matures along the secretory pathway, where it is subjected to glycosylation, phosphorylation and propeptide scission. Glycosylation is initiated in the ER and it continues in the Golgi apparatus, where glycosylation is refined to its mature state. N-Cadherin phosphorylation is essential for its interaction with β-Catenin and although the exact compartment in which this phosphorylation takes place is unknown, it is thought to occur prior to entry into the TGN (McEwen et al, 2014). In addition, Pro-N-Cadherin contains a Furin cleavage site, which is acted on in the TGN or in the space between this compartment and the cell surface (Nakayama, 1997). The data obtained here demonstrate that β-Catenin is required for efficient N-Cadherin delivery to the AC of NSCs and to the intercellular junctions of other cells. To determine if β-Catenin is implicated in the maturation of Pro-N-Cadherin, an antibody against the propeptide region of chicken Pro-N-Cadherin was developed (anti Pro-N-Cadherin), as well as a mutant form of N-Cadherin in which the Furin cleavage site was substituted by a Factor Xa cleavage consensus motive (FXa-N-Cadherin) to produce ^FXa^Pro-N-Cadherin, a variant that was not processed into mature N-Cadherin (Koch et al, 2004). Furthermore, we used an antibody that recognized the mature extracellular domain of chicken N-Cadherin (anti N-Cadherin: Fig. 6A). Interestingly, in Western blots of untransfected HH16 chicken neural tubes the Pro-N-Cadherin antibody detected two bands that migrated slower than the mature N-Cadherin (Fig. 6B). Thus, we transfected HH12 chicken neural tubes with N-Cadherin·ST to study the consequences of β-Catenin knockdown or over-expression on N-Cadherin processing, calculating the ratio between the Pro-N-Cadherin (upper band) and mature N-Cadherin.

**Figure 6.**
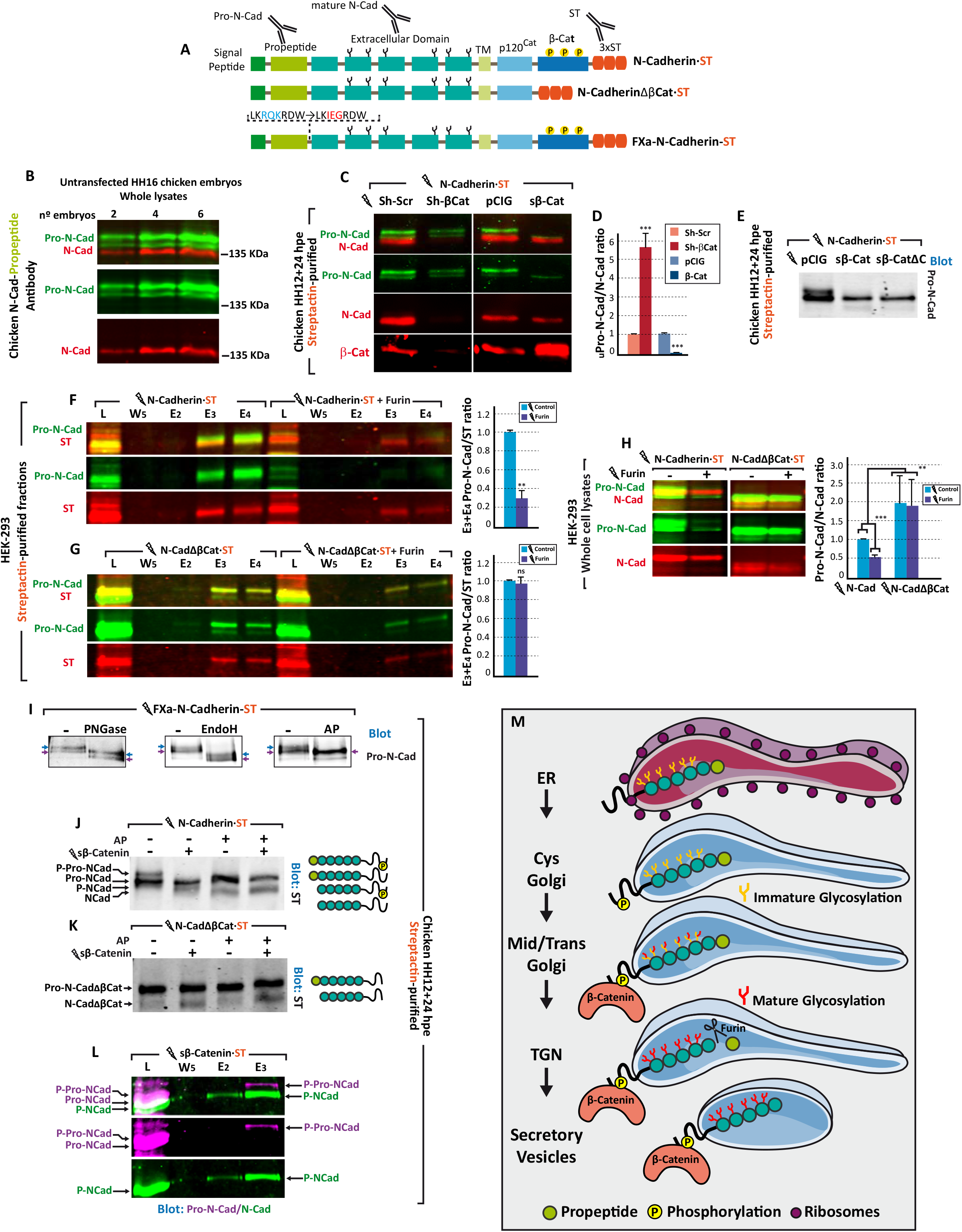
Pro-N-Cadherin needs to interact with β-Catenin to excise the propeptide but not for glycosylation. **(A)** Scheme of the N-Cadherin constructs and the regions of N-Cadherin targeted by the antibodies used. **(B)** Western blot of whole lysates from untransfected HH16 chicken embryos probed with antibodies against Pro-N-Cadherin (Green) and mature-N-Cadherin (Red). **(C)** Western blot of Streptactin purified N-Cadherin-ST electroporated into HH12 chicken neural tubes with Sh-βCat or sβ-Catenin. The blot was probed with antibodies against Pro-N-Cadherin (Green), Mature-N-Cadherin (Red) and β-Catenin (Red). **(D)** Bar-graph showing the control-normalized ratios between Pro-N-Cadherin and mature-N-Cadherin for each transfection. **(E)** As in C but using sβCatΔC, a sβ-Catenin mutant with no transcriptional activity. **(F)** Western blot showing Streptactin-purified fractions of Hek-293 cultures transfected with the Furin or control vector and N-Cadherin·ST. The blot was probed with antibodies against Pro-N-Cadherin (Green) and ST (Red): L, Lysate; W, Wash; E, Elution. The bar graph shows the control-normalized ratios between Pro-N-Cadherin and mature-N-Cadherin. **(G)** As in F but with N-CadherinΔβCat. **(H)** Western blot of whole cell lysates from an experiment similar to F and G but probed with antibodies against Pro-N-Cadherin (Green) and mature-N-Cadherin (Red) rather than ST. The bar graph shows the control normalized ratios between Pro-N-Cadherin and mature-N-Cadherin. Note that N-CadherinΔβCat is not converted into mature N-Cadherin by Furin. **(I)** Western blot of Streptactin purified N-Cadherin-ST electroporated into HH12 chicken neural tubes and treated with PNGase (degrades all N-linked glycosylation), EndoH (degrades immature N-linked glycosylation) or AP (Alkaline Phosphatase). The blot was probed with antibodies against Pro-N-Cadherin. **(J)** Blot of Streptactin purified fractions from HH12 chicken neural tubes transfected with N-Cadherin-ST, with or without sβ-Catenin, and treated with AP. A scheme of the proposed molecular nature of each band is shown to the right of the blot. **(K)** Same experiment as in J but with N-CadherinΔβCat. **(L)** Western blot of Streptactin purified fractions from HH12 chicken neural tubes transfected with sβ-Catenin·ST, and probed with antibodies against Pro-N-Cadherin (Purple) and mature-N-Cadherin (Green). Note that only the upper band of Pro-N-Cadherin binds to sβ-Catenin·ST. **(M)** Model displaying the sequence of events occurring in the course of N-Cadherin maturation: ER, endoplasmic reticulum; TGN, trans Golgi network. All the bar graphs shown in this figure represent the mean ± SD: ns, not significant; \**p* <0.05; \*\**p* <0.01; \*\*\**p* <0.001. Lightning bolts denote transfection.

Notably, β-Catenin knockdown significantly decreased the mature N-Cadherin relative to the Pro-N-Cadherin, whereas sβ-Catenin expression had exactly the opposite effect (Fig. 6C,D). From this experiment we also noticed two important things, that sβ-Catenin expression mainly reduced the upper band of Pro-N-Cadherin and that this effect was also obtained with sβ-CateninΔC, a mutant sβ-Catenin with no transcriptional activity (Fig. 5 and 6).

The ST-Streptactin purification system was then used on HEK-293 cells to study whether β-Catenin binding was required for Pro-N-Cadherin to be processed by Furin. Notably, Furin transfection significantly enhanced N-Cadherin propeptide cleavage (Fig. 6F) but not that of N-CadherinΔβCat (Fig. 6G). Accordingly, this deficient N-CadherinΔβCat propeptide cleavage provoked an increase in the Pro-N-Cadherin/N-Cadherin ratio (Fig. 6H). These results demonstrate that β-Catenin binding to N-Cadherin is required for propeptide cleavage and they strongly suggest that deficient propeptide cleavage prevents the maturation of N-CadherinΔβCat.

Glycosylation and phosphorylation are two post-translational modifications that may alter the molecular weight of N-Cadherin and that could therefore account for the shift in Pro-N-Cadherin mobility induced by sβ-Catenin. Thus, to eliminate one of the variables in this equation, we used the FXa-N-Cadherin·ST construct in which the Furin site is substituted with a FXa site (Fig. 6A). Like the endogenous Pro-N-Cadherin, the ^FXa^Pro-N-Cadherin generated by the FXa-N-Cadherin·ST construct was detected as a doublet by the Pro-N-Cadherin antibody when it was expressed in HH12 chick embryo neural tubes and purified on Streptactin columns (Fig. 6I). To define the molecular nature of the two Pro-N-Cadherin bands, we treated the purified ^FXa^Pro-N-Cadherin·ST with PNGase, a glycosidase that strips immature and mature glycosylation, with EndoH that only strips immature glycosylation, or with Alkaline Phosphatase (AP) that produces dephosphorylation. The two glycosidases induced a comparable band-shift of both ^FXa^Pro-N-Cadherin bands, indicating that both proteins were similarly glycosylated with immature glycosylation. Conversely, AP treatment converted the upper band into the lower one, demonstrating that the difference in molecular weight between the two ^FXa^Pro-N-Cadherin protein bands was due to their phosphorylation.

In a similar experiment we transfected N-Cadherin-ST in the presence or absence of sβ-Catenin, and we assessed the effect of AP on the purified fractions (Fig. 6J). The combined knowledge gained in these two experiments demonstrated that overexpressed N-Cadherin accumulated mainly as phosphorylated and unphosphorylated Pro-N-Cadherin, while sβ-Catenin promoted the conversion of phospho-Pro-N-Cadherin into phospho-N-Cadherin (Fig. 6J), consistent with the fact that β-Catenin binds mainly to phosphorylated Cadherins (McEwen et al, 2014; Stappert & Kemler, 1994). By contrast, transfection of N-CadherinΔβCat predominantly produced a single band that corresponded to the unphosphorylated Pro-N-CadherinΔβCat, and that was not affected by sβ-Catenin co-expression, or by AP. Hence, the effect of sβ-Catenin on the cleavage of the Pro-N-Cadherin propeptide required direct N-Cadherin/β-Catenin binding and like E-Cadherin (McEwen et al, 2014; Stappert & Kemler, 1994), N-Cadherin phosphorylation sites are located in the β-Catenin binding domain (Fig. 6K). To further support these data, we electroporated HH12 chicken neural tubes with sβ-Catenin·ST and purified it along with its associated endogenous proteins on Streptactin columns. Consistently, only the upper band of Pro-N-Cadherin and the mature N-Cadherin were co-purified with sβ-Catenin·ST (Fig. 6L). In summary, the data obtained demonstrate that the interaction of sβ-Catenin with the phosphorylated form of Pro-N-Cadherin in the TGN enables the excision of the propeptide and the delivery of N-Cadherin to the AC (Fig. 6M).

### The persistence of Pro-N-Cadherin induces the loss of apico-basal polarity, causing aberrant delamination, a breach of the basal membrane and mesenchymal invasion

EMT is a process that involves a loss of cell adhesion and thus, one effect of the EMT genes induced (e.g., Snail) is the inhibition of E-Cadherin transcription. Indeed, the non-adhesive N-cadherin precursor is expressed abundantly at the surface of invasive melanoma and brain tumour cells, the invasive potential of which is directly correlated to the ratio between Pro-N-Cadherin and mature N-Cadherin (Maret et al, 2010). Thus, we wondered whether the persistence of Pro-N-Cadherin would affect the polarity and cohesion of NSCs in the neuroepithelium. In chicken embryonic NSCs, over-expressed N-Cadherin-GFP accumulated at the ACs, and its additional accumulation in other compartments of the cell probably reflects a delay in its maturation (Fig. 4E). In fact, Pro-N-Cadherin was abundant around the nucleus in NSCs transfected with N-Cadherin, yet abundant mature N-Cadherin was also found throughout the cell, mostly apical to the nucleus, in areas where mature and immature N-Cadherin coexisted (Fig. 7A,A’ see arrows). After delamination, both Pro and mature N-Cadherin accumulated in the immediacy of the nucleus and in the newly forming neurites (Fig. 7A,A’,A”; see arrowheads), reflecting the reported Golgi relocation that occurs during this process (Taverna et al, 2016). In accordance with the immunostaining, Streptactin purification revealed that a significant proportion of the transfected N-Cadherin·ST remained as Pro-N-Cadherin (Fig. 7B), even though these neural tubes did not display any aberrant growth or altered apico-basal polarity. Conversely, abundant ^FXa^Pro-N-Cadherin but not mature ^FXa^N-Cadherin was detected around the nucleus of the cells transfected with FXa-N-Cadherin. Notably, all the transfected cells had lost their apical/basal polarity and although some of them invaded the ventricle (Fig. 7C,C’), most of these cells accumulated in the mantle zone, distorting the normal neural tube structure (Fig 7C,C”;). Indeed, mature ^FXa^N-Cadherin was not purified from FXa-N-Cadherin·ST transfected neural tubes (Fig. 7D).

**Figure 7.**
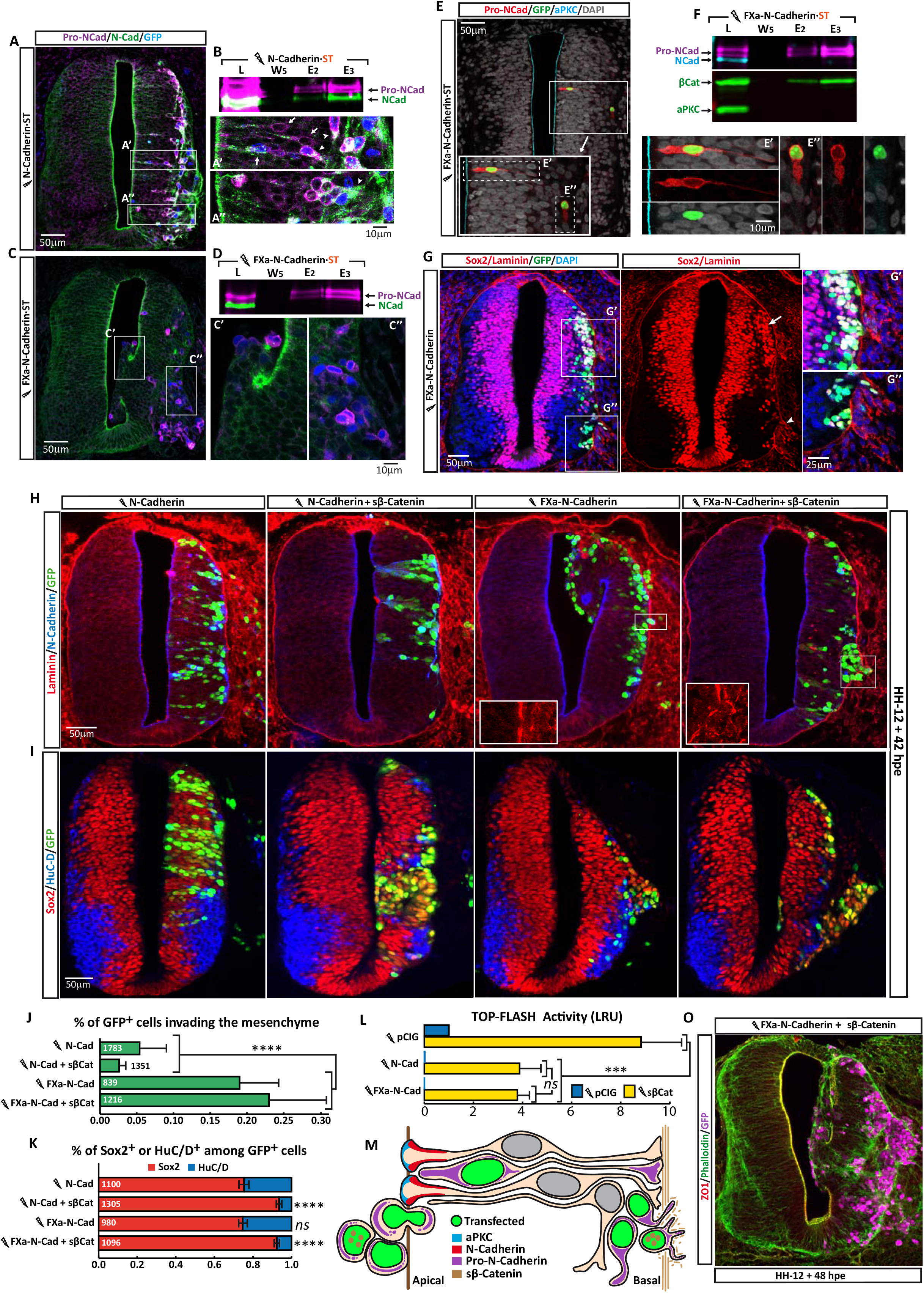
Pro-N-Cadherin persistence induces a loss of apico-basal polarity, causing aberrant delamination, ventricle invasion and basal membrane rupture. **(A)** HH12 chicken neural tubes 48 hpe with N-Cadherin, and probed with antibodies against mature-N-Cadherin (Green) and Pro-N-Cadherin (Purple), and with nuclear GFP indicating transfection (Blue). The areas labelled as A’ and A”; are amplified on the right, and the arrows indicate neural precursors while the arrowheads show the delaminated neurons. **(B)** Western blot of Streptactin-purified fractions of HH12 chicken neural tubes transfected with N-Cadherin·ST probed with antibodies against mature-N-Cadherin (Green) and Pro-N-Cadherin (Purple): L, Lysate; W, Wash; E, Elution. **(C,D)** A similar experiment to A,B but using Pro-N-Cadherin·ST. **(E)** Weakly transfected HH12 chicken neural tubes with Pro-N-Cadherin probed with antibodies against Pro-N-Cadherin (Red) and aPKCι (Magenta), stained with DAPI (Grey), and with nuclear GFP expression indicating transfection (Green). The areas labelled as E’ and E”; are amplified on the right, and the channels were split for clarity. **(F)** Western blot of Streptactin-purified fractions from HH12 chicken neural tubes transfected with Pro-N-Cadherin·ST and probed with antibodies against Pro-N-Cadherin (Purple), aPKCι (Magenta), mature-N-Cadherin (Green) and β-Catenin (Green). Note that Pro-N-Cadherin·ST binds to β-Catenin but not aPKCι. **(G)** HH12 chicken neural tubes 48 hpe with Pro-N-Cadherin stained with antibodies against Sox2 (Red) and Laminin (Red), and with DAPI (Blue). Nuclear GFP indicates transfection (Green). The areas labelled as G’ and G”; are amplified on the right. **(H,J)** Images and quantification of HH12 chicken neural tubes at 42 hpe with N-Cadherin or Pro-N-Cadherin, with or without sβ-Catenin, and probed with antibodies against Laminin (Red) and N-Cadherin (Blue), with GFP (Green) indicating transfection. The amplified areas in the Pro-N-Cadherin and Pro-N-Cadherin plus sβ-Catenin panels show details of basal membrane breaching. The bar graph represents the mean ± SD of the cells invading the mesenchyme in each treatment. **(I,K)** As in H,J but stained with antibodies against Sox2 (Red) and HuC/D (Blue), and GFP (Green) indicating transfection. The bar graph represents the mean ± SEM of the Sox2+ and HuC/D+ cells among the transfected population. **(L)** The bar graph represents the mean ± SD (three independent experiments) of a Luciferase reporter assays using TOP-FLASH vector 48 hpe with pCIG, N-Cadherin or Pro-N-Cadherin, co-transfected with sβ-Catenin. **(M,O)** Scheme and a representative image showing the growth aberrations in a HH12 chicken neural tube 48 hpe with Pro-N-Cadherin and sβ-Catenin, stained with an antibody against ZO1 (Red), and for Actin (Phalloidin, Green) and GFP (Magenta, indicating transfection). At least 5 embryos were used per treatment in the bar graphs and the total number of cells counted in each case is indicated within the bars: *ns*, not significant; \*\*\**p* <0.001. Lightning bolts denote transfection.

To better understand the effect of impairing Pro-N-Cadherin maturation, we assessed slices with mild transfection to visualize isolated FXa-N-Cadherin transfected cells (Fig. 7E). In the cells located in the proliferative zone, ^FXa^Pro-N-Cadherin accumulated in aberrant apical processes that did not form AJs and did not reach the aPKC layer (Fig 7E’). Conversely, the delaminated cells expressing ^FXa^Pro-N-Cadherin in the mantel zone remained as unpolarised cells emitting short cytoplasmic processes (Fig. 7C,E”;). Consistent with its incapacity to form AJs, purification of ^FXa^Pro-N-Cadherin·ST revealed that while it associated with β-Catenin, the complex did not contain aPKC (Fig. 7F). Notably, the groups of cells expressing ^EXa^Pro-N-Cadherin that accumulated in the mantel zone had a generous proportion of atopic Sox2+ progenitors (Fig. 7G). Moreover, the ^FXa^Pro-N-Cadherin expressing clumps pressured the basal membrane, distorting it and even breaching it at certain debilitated sites (Fig. 7G,G’,G”;). As the presence of Sox2+ cells in the mantel zone could be due to premature delamination of progenitors or to their inhibited differentiation, we electroporated HH12 chicken embryo neural tubes with vectors expressing N-Cadherin or ^FXa^Pro-N-Cadherin, and 42 hpe we studied the proportion of progenitors (Sox2+) and neurons (HuC/D+) among the transfected cells. Although transfection of FXa-N-Cadherin severely affected the distribution of the transfected cells in the neural tube, it did not alter the proportion of progenitors and neurons (Fig. EV3 A-D).

We previously showed that sβ-Catenin maintains stemness in NSCs (Herrera et al, 2014). Moreover, E-cadherin can prevent cell growth and transformation by recruiting β-Catenin to the AJs (Gottardi et al, 2001; Stockinger et al, 2001). Thus, we wondered whether the invasion of the mesenchyme caused by FXa-N-Cadherin expression was due to deficient buffering of β-Catenin transcriptional activity. However, N-Cadherin and FXa-N-Cadherin reduced sβ-Catenin transcriptional activity in a very similar way (Fig. 7L) and in fact, sβ-Catenin did not affect the total number of cells that invaded the mesenchyme in response to FXa-N-Cadherin (Fig. 7H,J). Notably however, sβ-Catenin significantly increased the proportion Sox2+ NSCs among the cells that invaded the mesenchyme in response to FXa-N-Cadherin expression (Fig. 7IK, see also the scheme and a representative image of the invasive phenotype induced by FXa-N-Cadherin in Fig. 7M,O).

In summary, a failure in N-Cadherin maturation disrupts the formation of AJs, on the one hand weakening the apical surface of the neuroepithelium and provoking invasion of the ventricle, and on the other triggering the premature delamination of neural precursors that accumulate in the mantel zone, invading the surrounding mesenchyme. In these circumstances, although the oncogenic β-Catenin does not initiate invasion it maintains the stemness of the invading cells, thereby enhancing their oncogenic potential. Consequently, mutations in N-Cadherin or in other genes that participate in N-Cadherin processing, will abolish the onco-suppressor activity of sβ-Catenin, causing sβ-Catenin dependent tumours to progress.

## DISCUSSION

### β-Catenin binding promotes Pro-N-Cadherin processing and transport to the AC

We previously demonstrated that stable forms of β-Catenin (sβ-Catenin) induce oversized ACs containing abnormally large amounts of N-Cadherin, β-Catenin, actin and active aPKC (Herrera et al, 2014). We showed that sβ-Catenin induces both the transcription and the recruitment of aPKC to the ACs, and we demonstrated that the hyperpolarization induced by sβ-Catenin was reverted by a dominant negative form of aPKCι. Moreover, we found that the depolarization of the neuroepithelium induced by aPKMz, an active form of aPKC, could be reverted by sβ-Catenin, which concentrated all the aPKC activity in the AC and prevented its ectopic activation in the basolateral compartment. Accordingly, we wondered whether N-Cadherin, β-Catenin and aPKC themselves formed a complex and if so, whether this complex was formed at the AC or whether it was pre-assembled and transported there.

We demonstrate here that aPKC co-localizes with N-Cadherin or β-Catenin exclusively at the AC of NSCs, whereas N-Cadherin also co-localizes with β-Catenin in the Golgi apparatus, albeit less intensely than in the AC. Nevertheless, the exact point at which the Cadherin/β-Catenin complex is initially established remains unclear. It was first demonstrated that β-Catenin binds to E-Cadherin early in the biosynthetic pathway (Hinck et al, 1994) and later, it was proposed that the E-Cadherin/β-Catenin complex was formed in the ER. Indeed, the formation of this complex allowed the efficient ER exit and basolateral membrane targeting of a chimeric protein that contains the cytoplasmic domain of E-Cadherin in polarized MDCK cells (Chen et al, 1999). In addition, β-catenin was seen to bind to PX-RICS, a GTPase-activating protein (GAP) for Cdc42 that associates to dynein-dynactin, promoting ER-to-Golgi transport of the N-cadherin/β-catenin complex. Indeed, the accumulation of the N-cadherin/β-catenin complex in the ER as a result of PX-RICS downregulation leads to a decrease in cell adhesion (Nakamura et al, 2010; Nakamura et al, 2008). Alternatively, in vitro studies demonstrated that β-Catenin presents an ~800-fold higher affinity for E-Cadherin when this is phosphorylated (McEwen et al, 2014) and that the β-Catenin/E-Cadherin complex forms before the excision of the propeptide domain of E-Cadherin (Wahl et al, 2003). Therefore, we assume that Pro-Cadherins are phosphorylated before binding tightly to β-catenin and indeed, we found that β-catenin can be purified exclusively with the phosphorylated form of Pro-N-Cadherin.

Although the compartment in which Pro-Cadherins are phosphorylated is unknown (McEwen et al, 2014) the phosphorylated band of Pro-N-Cadherin can be effectively deglycosylated by EndoH, an endoglycosidase that only acts on high-mannose immature glycosylation (found in the ER and cys-Golgi apparatus). Hence, Pro-N-Cadherin phosphorylation must take place prior to entry into the medial/trans-Golgi compartment. Here we show that N-Cadherin and β-Catenin co-localize more in the post-Golgi space than in the Golgi apparatus itself, reaching a maximum at the AC. Thus, it is possible that the phosphorylated form of Pro-N-Cadherin and β-Catenin assemble in the Golgi. Interestingly though, we observed that N-CadherinΔβ-Cat, a mutant N-Cadherin that lacks the β-Catenin binding domain and consequently, its main phosphorylation sites, does not accumulate in the ER but rather, it accumulates as Pro-N-Cadherin in Golgin97 vesicles in the TGN.

E-Cadherin mutants that cannot bind to β-Catenin have consistently been reported to accumulate in intracellular compartments, including the TGN (Miyashita & Ozawa, 2007). Golgin97 is an essential component of the carriers transporting E-Cadherin out of the TGN into Rab11-positive recycling endosomes on its way to the cell surface (Lock et al, 2005; Lock & Stow, 2005). Hence, although the Pro-Cadherin/β-Catenin complex may already be formed in the ER, their interaction remains weak until Pro-N-Cadherin is phosphorylated somewhere before the mid-Golgi stage, thereafter forming a stable complex with β-Catenin. Likewise, Pro-N-Cadherin does not need to be coupled to β-Catenin to leave the ER or to reach the TGN but conversely, the lack of β-catenin prevents the excision of the propeptide and causes Pro-N-Cadherin to accumulate in the TGN. Thus, β-catenin binding to Pro-N-Cadherin seems to be especially critical for the propeptide excision step in the TGN and consequently, for the formation of functional apical AJs. By contrast, co-localization of aPKC with either N-Cadherin or β-Catenin occurs almost exclusively at the AC. Our results demonstrate that N-Cadherin/β-Catenin complexes and aPKC are delivered separately to the AC, yet no apical aPKC accumulation was observed in the absence of N-Cadherin/β-Catenin. On reaching the apical membrane, aPKC forms a complex with Par3 and Par6, where the combined activity of Cdc42 and PDK1 activates the kinase activity of aPKC, which like β-Catenin and N-Cadherin is required to maintain apico-basal polarity (Chen & Zhang, 2013). Therefore, although more evidence will be required to fully demonstrate the model, we speculate that the boundary region between the apical and sub-apical domains where the three molecules co-exist serves as a docking station where β-Catenin interacts with Par3 through its PDZ binding domain (Gujral et al, 2013), capturing Par3/Par6/aPKC complexes that are then shunted to the apical membrane.

### The activity of sβ-Catenin as a tumour suppressor depends on its capacity to promote N-Cadherin processing

WNT type Medulloblastoma is mainly caused by stable forms of β-catenin (sβ-catenin) and it is the least aggressive of the four types of medulloblastomas, mainly due to its low invasiveness and the fact that it rarely metastasizes. Notably, the expression of sβ-catenin significantly increases survival in a mouse model of Shh-type medulloblastoma (Poschl et al, 2014), which indicates that sβ-catenin acts simultaneously as both an oncogene and a tumour suppressor. Among the multiple changes that occur during the development of invasive malignancies (Hanahan & Weinberg, 2000), two major events are the onset of uncontrolled cell proliferation and the loss of cell adhesion, together driving the invasion of local tissues (Birchmeier et al, 1996). E-Cadherin is considered a suppressor of invasion because it increases cell adhesion in epithelial tissues (Gumbiner, 2000; Larue et al, 1996) but also, because it binds to β-Catenin and prevents its transforming activity (Gottardi et al, 2001; Jeanes et al, 2008). Indeed, epithelia that express E-Cadherin strongly are more resistant to being transformed by oncogenic mutations of β-Catenin (Huels et al, 2015).

We previously reported that the activity of sβ-catenin at AJs maintains NSCs as progenitors but it also enhances their polarization and adhesiveness, which in turn limits their proliferation and migration (Herrera et al, 2014). We now propose that the function of sβ-catenin as a tumour suppressor depends on its ability to promote the maturation of Pro-N-cadherin, with the consequent formation of larger AJs. This effect not only strengthens cell adhesion but also, it restricts the delamination and dissemination of sβ-catenin cells. Conversely, mutations that alter Pro-N-Cadherin maturation would prevent the formation of functional AJs, causing a loss of cell adherence and as a result, an increase in the cell’s invasive potential. A loss of adhesion is also commonly associated to an increase in apoptosis (Carmeliet et al, 1999) due to the activation of the Fas death receptor (Gagnoux-Palacios et al, 2018). However, such a preventive apoptotic response is often lost in tumours due to their oncogenic activation, making a loss of adhesion particularly damaging in such environments (Gagnoux-Palacios et al, 2018).

We show here that mutations that prevent Pro-N-Cadherin processing induce a loss of polarity, ventricle invasion, premature delamination, ectopic progenitor accumulation in the mantle zone and a breach of the basal membrane. Notably, although we found that Pro-N-Cadherin still binds to β-Catenin, the combination of Pro-N-Cadherin and sβ-Catenin prevented cell differentiation. Moreover, while sβ-Catenin transcriptional activity does not seem to participate in the initial events of mesenchyme invasion, the fact that the invading cells remain as proliferating NSCs greatly increases their oncogenic potential. Notably, CD133 and Sox2 expressing NSCs are thought to be the cell populations propagating medulloblastoma (Parada et al, 2017). It is significant that a cytoplasmic (perinuclear) rather than cell-surface distribution of E-cadherin has often been reported in carcinoma cells, suggesting that the transport of E-cadherin to the cell surface is impaired in some carcinomas (Carpenter et al, 2002). Moreover, the expression of surface Pro-N-cadherin in melanoma and glioblastoma cell lines promotes tumour cell invasion, and a positive ratio of Pro-N-Cadherin/N-Cadherin correlates with increased malignancy in different human cancers (Maret et al, 2010). In addition, Pro-N-cadherin has been detected on the cell surface of a highly invasive sub-population of triple-negative breast tumour cells (Nelson et al, 2016). Together these data indicate that the loss of adhesiveness caused by Pro-N-Cadherin at the cell surface may have an accumulative effect on the oncogenic capacity of other oncogenes, as occurs with sβ-Catenin. Therefore, it would be very interesting to check whether cytoplasmic or cell surface Pro-N-Cadherin can be observed in the medulloblastomas that metastasize or progress to a higher grade malignancy.

## METHODS

### Commercial antibodies and chemicals

Mouse antibodies against HuC/D (Molecular Probes #A21271; 1:500), β-Catenin (Sigma #C7207, 1:200), β-Catenin (Cell signaling, #9562; 1:500), Golgin97 (Invitrogen #A21270; 1:100), Strep-tag (IBA #2-1507-001; 1:500), CLIMP63 (US Biological #C5840-93; 1:1000), aPKCζ/λ (Santa Cruz #SC-17781; 1:200) and GM130 (BD Biosciences #610822; 1:500).

Rat antibodies against N-Cadherin (Invitrogen #13-2100; 1:200)

Rabbit antisera against Laminin (Sigma #L9393; 1:1000), Sox2 (Invitrogen #48-1400; 1:500) and Calreticulin (Abcam #ab2907; 1:1000).

Chemicals: Rhodamine-Phalloidin (Invitrogen, #R415; 1:250).

### In house antisera

The rabbit antisera used to detect FLAG and RFP were described in (Herrera et al, 2014), while the Pro-N-Cadherin antiserum was raised against a GST (Glutathione S-Transferase) fusion protein with the pro-peptide region of chicken N-Cadherin (aa 29-164).

### DNA constructs

Proteins were expressed using the pCIG, pCIEGO, pIRES or pCS2 vectors and short hairpin inhibitory RNAs were generated with pSHIN. To generate ST-tagged proteins, three copies of the *Strep-tag^®^* II peptide (WSHPQFEK) were cloned at the C-terminus of the proteins indicated. Deletion and substitution mutants were generated through standard techniques and all constructs were sequenced prior to use.

#### Vectors

pCIG (Chicken β-Actin Promoter/CMV-IE enhancer, EGFP expressing bicistronic vector), pCIEGO (pCIG with no IRES-EGFP), pIRES (bicistronic Human CMV-IE promoter/enhancer, EGFP expressing vector), pCS2 (Simian CMV-IE94 promoter) and pSHIN (expresses Short Hairpin inhibitory RNAs driven by the human H1 promoter and EGFP driven by the artificial SRα promoter).

#### pCIG constructs

Chicken N-Cadherin; N-Cadherin·ST, ST tagged Chicken N-Cadherin; N-CadherinΔβCat, Chicken N-CadherinΔ844-912; N-CadherinΔβCat·ST, ST tagged Chicken N-CadherinΔ844-912; XFa-N-Cadherin, the Furin consensus cleavage sequence (LKRQKRDW) amino acids 159-166 of Chicken N-Cadherin were substituted by the Factor Xa cleavage sequence (LKIEGRDW); FXa-N-Cadherin·ST, ST tagged Chicken FXa-N-Cadherin; sβ-Catenin, Human β-Catenin-S33Y; sβ-Catenin·ST, ST tagged Human β-Catenin-S33Y; sβ-CateninΔC, Human β-Catenin-S33Y-Δ674-782; aPKCι·ST, ST tagged Human aPKCι; TCF3·VP16, in which the HMG box of mouse TCF3 was fused to the VP16 transcriptional activator.

#### pCIEGO construct

sβ-Catenin, Human β-Catenin-S33Y.

pIRES construct: Human Furin.

#### pCS2 constructs

N-Cadherin·GFP, Chicken N-Cadherin with EGFP fused to the C-terminus; mbGFP, green fluorescent protein with glycosylphosphatidylinositol fused to the C-terminus; mbRFP, red fluorescent protein with glycosylphosphatidylinositol fused to the C-terminus.

#### pSHIN constructs

Sh-Scr, Scrambled ShRNAi sequence (CCGGTCTCGACGGTCGAGT); Sh-βCat, ShRNAi targeting chicken β-Catenin (ATCCCAGAACTGACCAAAC).

### Chick embryo *in ovo* electroporation

Eggs from White-Leghorn chickens were incubated at 37.5 °C in an atmosphere of 45% humidity and the embryos were staged according to Hamburger and Hamilton (HH: (Hamburger & Hamilton, 1992). Chick embryos were electroporated with column purified plasmid DNA (1-2 μg/μl) in H_2_O containing Fast Green (0.5 μg/μl). Briefly, plasmid DNA was injected into the lumen of HH12 neural tubes, electrodes were placed on either side of the neural tube and electroporation was carried out by applying five 50 ms square pulses using an Intracel Dual Pulse (TSS10) electroporator set at 25 V. Transfected embryos were allowed to develop to the specific stages and then dissected under a fluorescence dissection microscope.

### Purification of Strep-tag II tagged proteins on Strep-Tactin columns

*Strep-tag*^®^ II is a short peptide (8 amino acids, WSHPQFEK) that binds with high affinity/selectivity to Strep-Tactin^®^, an engineered streptavidin. To tag proteins with Strep-tag, three copies of the Strep-tag peptide were inserted at the C-terminus of the selected proteins. For purification experiments, chicken embryos or cell cultures electroporated with tagged proteins were dissolved in PIK buffer (Tris-HCl 20 mM [pH 7.4], NaCl 137 mM, Glycerol 10%, NP-40 1%, CaCl2 1 mM, MgCl_2_ 1 mM, Aprotinin 10 μg/ml, Leupeptin 10 μg/ml and PMSF 1 mM), with 1 ml of PIK buffer used for each 10 cm culture dish or 15 embryos (HH12 embryos, 24 hpe). For co-purification experiments, GFP+ areas of electroporated neural tubes were dissected out in cold PBS and then the protein complexes were mildly cross-linked with 1% formaldehyde for 10 minutes at room temperature. The cross-linking reaction was stopped with glycine up to 125 mM and the cross-linked embryos were then dissolved by sonication (2 × 30 seconds at 10 μm amplitude) in ice-cold SDS lysis buffer (1% SDS, 10 mM EDTA, 50 mM Tris-HCl [pH 8.1], Aprotinin 10 μg/ml, Leupeptin 10 μg/ml, PMSF 1 mM, NaF 5 mM, NaVa 1 mM). The protein samples were diluted in a dilution buffer (1.1% Triton X-100, 1.2 mM EDTA, 17 mM Tris-HCl [pH 8.1], 170 mM NaCl, Aprotinin 10 μg/ml, Leupeptin 10 μg/ml, PMSF 1 mM, NaF 5 mM, NaVa 1 mM) to a final concentration of 0.1% SDS, 0.1% Triton X-100, 2 mM EDTA, 20 mM Tris-HCl [pH 8.1], 150 mM NaCl, Aprotinin 10 μg/ml, Leupeptin 10 μg/ml, PMSF 1 mM, NaF 5 mM, NaVa 1 mM. In these experiments the insoluble material was removed by centrifugation at 18,500 × g for 5 minutes. Purification was performed on prepacked Strep-Tactin^®^ (0.2 ml bed volume (bv) gravity-SuperFlow^®^ columns: IBA Lifesciences, Goettingen, #2-1209-550) following the manufacturer’s instructions and using the reagents provided with the columns, except that up to 1% of Triton X-100 (purifications) or 1% of Triton X-100 and 0.1% SDS (co-purifications) was added to the wash and elution buffers. Briefly, columns were pre-stabilized with 5 bvs of lysis buffer and 1 ml of lysate was applied to each 0.2 ml column, washed with 5-15 bvs, and finally eluted in 100 μl aliquots. Purified proteins were stored at −80 for further use.

### Immunoblotting

Purified fractions were mixed with 5x SDS Laemmli sample buffer (1x = 2% SDS, 65 mM Tris-HCl [PH 6.8], 10% Glycerol, 100 mM DTT and Bromophenol Blue 0.5 mg/ml), boiled for 5 mins (or 20 mins for cross-linked samples), resolved by SDS-PAGE (8% gels) and transferred to nitrocellulose membranes. The membranes were blocked with Odyssey Blocking Buffer (OBB^®^), incubated with the primary antibodies in OBB containing 0.2% of Tween-20, washed 3 times with TTBS (150 mM NaCl, 0.1% Tween-20 and 20 mM Tris-HCl [pH 7.4]) and incubated with Alexa labelled secondary antibodies in OBB containing 0.2% of Tween-20 and 0.01% of SDS. After three final washes in TTBS, the membranes were allowed to dry and the fluorescence was detected on the Odyssey Infrared Imaging System (LI-COR). The expression values were quantified with ImageJ and the molecular weights were calculated using Bio-Rad Precision Molecular Weight Markers. The values shown represent the means ± SD of three independent experiments.

### Immunohistochemistry

Embryos were fixed for 4 hours (to HH 20) or overnight (beyond HH 20) at 4 °C in 4% paraformaldehyde and immunostaining was performed on vibratome sections (40 μm) following standard procedures. After washing in PBS-0.1% Triton X-100, the sections were incubated with the appropriate primary antibodies and developed with Alexa or Cyanine conjugated secondary antibodies. After staining, the sections were mounted and examined on a Leica SP5 or a Zeiss Lsm 780 multiphoton microscope.

### HEK293 Cell fractionation

Confluent cultures of HEK-293 cells in 10 cm dishes were rinsed once with PBS (supplemented with 1 mM CaCl2 and 1 mM MgCl_2_), scraped-off in 1 ml of ice cold hypotonic buffer (1 mM sodium bicarbonate [pH 7.4], 10 μg/ml Aprotinin, 10 μg/ml Leupeptin and 1 mM PMSF) and homogenized with a glass-Teflon Potter-Elvehjem. The lysates were transferred to 1.5 ml tubes and centrifuged at 1000 × g for 10 min at 4 °C. The supernatant was decanted and the pellet homogenized again as described above. The two resultant supernatants were pooled and centrifuged at 25,000 × g for 30 min at 4 °C. The first and second pellets contained the internal membranes and the plasma membrane, respectively, and they were washed once with the same buffer and then resuspended in 1x Laemmli sample buffer.

### Glycosidase and Alkaline phosphatase treatment

Aliquots of Streptactin purified proteins were used to study phosphorylation (with AP) and glycosylation (with EndoH or PNGase). Briefly, the AP reaction was carried out at 37 °C for 1 h in a volume of 80 μl containing: 40 μl of Streptactin purified protein, 20 μg/ml Aprotinin, 20 μg/ml Leupeptin, 1 mM PMSF, 50 mM Tris-HCl [pH 9], 1 mM MgCl_2_ and 2 μl of CIAP (Calf intestine AP, 30 U/μl: Takara#2250A). For the Endo H reactions, 36 μl of purified protein was incubated at 100 °C for 10 min with denaturing buffer (0.5% SDS, 40 mM DTT) in a final volume of 40 μl, and the volume was then raised to 80 μl by adding sodium citrate [pH 5.5] up to 50 mM. Each sample was then split into two and incubated in the presence or absence of 2 μl of EndoH (50 U/μl: NEB#P0702S) for 1h at 37 °C. The PNGase-F reaction was performed as for Endo H but the reaction buffer contained 50 mM Sodium Phosphate [pH 7.5], 1% NP40 and PNGase-F (50 U/μl: NEB#P0704S).

### Calculation of the Mander’s correlation coefficients

To analyse co-localization, 8192×8192 pixel images were acquired on either a Leica SP5 or a Zeiss Lsm780 confocal microscope. The area of interest was outlined manually in each dual-labelled confocal slice and then processed with the Coloc2 plugin of the Fiji version of the ImageJ image processing software. To calculate the fraction of N-Cadherin that co-localized with aPKC, β-Catenin, GM130 or Golgin97, Mander’s coefficients (with thresholds: (Manders et al, 1993) and the associated Costes P-values (Costes et al, 2004) were obtained for every region. At least 10 different images were analysed for each coefficient and all the coefficients were verified with the Costes statistical significance test.

### Statistical Analyses

In Figures 2D, 2G, 4F, 5F&G, 5H, 6J, 6L and Supp. 3D, each experimental condition was compared with every other experimental condition using an 1-way ANOVA with a Tukey’s multiple comparisons test. For Figures 3B, 5D and 5H each experimental condition was compared with its control condition, and an unpaired t test was performed. For Figure 6K each experimental condition was compared with the control using an 1-way ANOVA with a Tukey’s multiple comparisons test. All statistical analyses were calculated using Graphpad Prism 6 software and significance was assumed when p < 0.05(*), p<0.01(**) and p<0.001(***).

## ACKNOWLEDGEMENTS

The authors are indebted to E. Rebollo for her invaluable technical assistance at the AFMU Facility (IBMB). We want to thank Dr Elisa Marti (IBMB, Barcelona) and Nabil G. Seidah (IRCM, Montreal) for providing DNAs. The monoclonal antibodies were obtained from the Developmental Studies Hybridoma Bank, developed under the auspices of the NICHD and maintained by The University of Iowa (Department of Biological Sciences, Iowa City, Iowa 52242). The work in S.P.’s laboratory was supported by grants BFU2014-53633-P and BFU2017-83562-P.

## AUTHOR CONTRIBUTIONS

A.H. conceived and performed most experiments, analysed the data and discussed the results. A.M. performed the experiments and provided technical support for all the experiments. B.T. adapted the Strep-tag^®^ purification system to chicken samples. S.P. conceived the experiments, analysed the data, discussed the results and wrote the manuscript.

## EXPANDED VIEW FIGURE LEGENDS

**Figure EV1.**
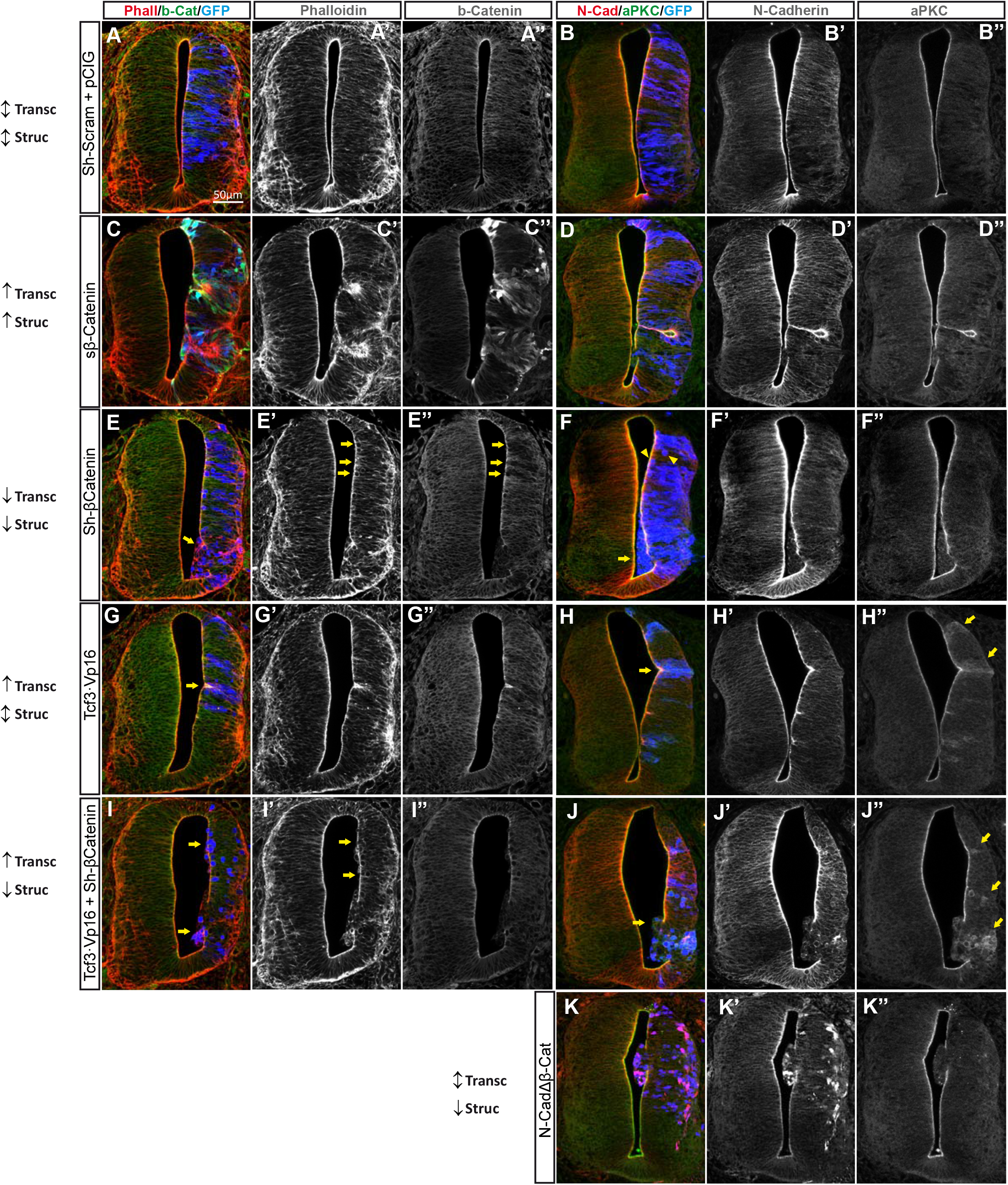
β-Catenin induces apical localization of aPKC through its binding to N -Cadherin. **(A)** HH12 Chicken neural tubes were transfected for 48 with different molecular tools intended to independently manipulate the transcriptional and structural activities of β-Catenin. The effect caused by each tool on the transcriptional and structural activities is indicated at the side of each row: ↑increased activity, ↓decreased activity, ↕ no change. In addition, pCIG stands for the empty vector, sβ-Catenin is an oncogenic form of β-Catenin (β-Catenin^S33Y^), Sh-Scram and Sh-βCat mean scrambled and anti β-Catenin Short Hairpin inhibitory RNAs, respectively. TCF3·VP16 is a constitutive activator of TCF dependent transcription and N-CadherinΔβCat is the N-Cadherin mutant lacking the β-Catenin binding domain. The figure shows representative images of the transfected neural tubes used for the quantification in Figure 3 and the channels are shown separately in grey scale for clarity. The slices were stained for F-Actin (Phalloidin, Red), β-Catenin (Green) and GFP (Blue, indicating transfection) in panels A,C,E,G and I, or for N-Cadherin (Red), aPKC (Green) and GFP (Blue, indicating transfection) in panels B,D,F,H,J, and K. The arrows in panels E, E’, E”; indicate Sh-βCat transfected cells with a very round morphology, some of them protruding into the ventricle. Knockdown of β-Catenin by Sh-βCat is clearly evident in panel E”;. In panels F, the arrowheads and arrow point to rounded and protruding cells, respectively. In panels G and H, the arrows pinpoint small invaginations caused by Tcf3·Vp16, while the arrows in panel H”; indicate the accumulation of cytoplasmic aPKC in the transfected areas. Arrows in panel I and I’ point to cells protruding into the ventricle and the arrows in panel J”; denote the accumulation of cytoplasmic aPKC in cells expressing a combination of Tcf3·Vp16 and Sh-βCat. Note that in this case the knockdown of β-Catenin caused a loss of apical-basal polarity rather than the typical small invaginations normally induced by Tcf3·Vp16.

**Figure EV2.**
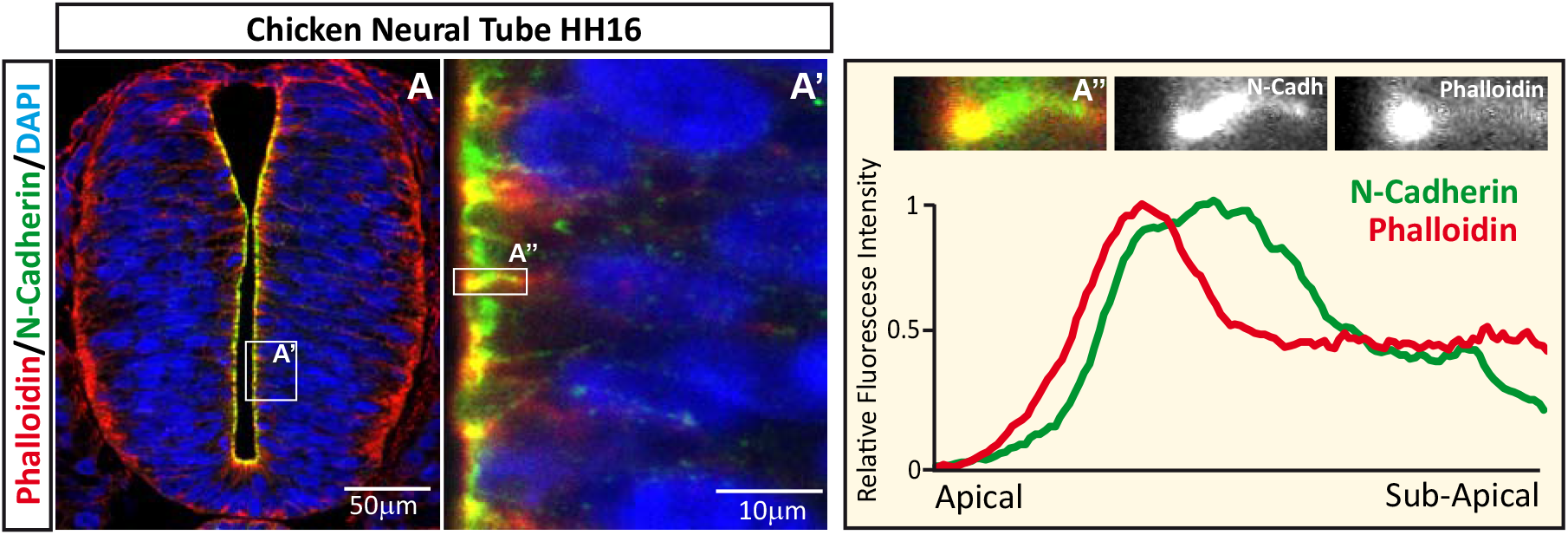
Actin expression can be detected apically and overlapping the N-Cadherin domain at the AC of NSCs. HH16 chicken neural tubes were stained for F-Actin (Phalloidin, Red), N-Cadherin (Green) and DAPI (blue). The area labelled as A’ is amplified in the right-hand panel, the plot shows the pixel intensity profile for N-Cadherin and Phalloidin in the area labelled as A”;. The area quantified is shown above the profile and the two channels are displayed separately in a grey scale for clarity.

**Figure EV3.**
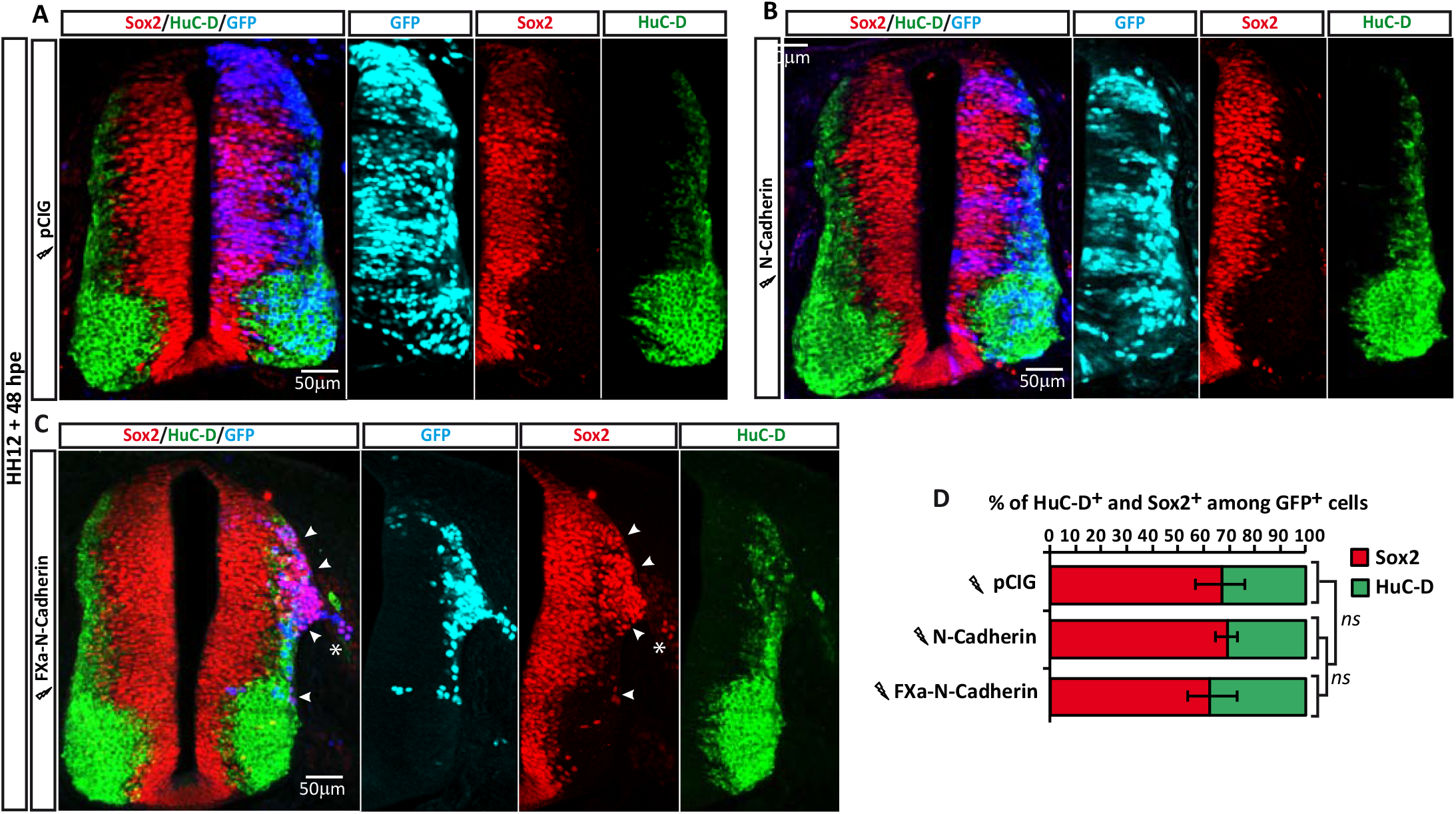
Pro-N-Cadherin expression does not change the proportion of Sox2+ and HuC/D+ cells among the NSCs. HH12 chicken neural tubes 48 hpe with control, N-Cadherin or Pro-N-Cadherin, and developed with Sox2 (Red) and HuC/D (Green). Nuclear GFP expression indicates transfection (Blue, or Magenta in the split channel composition). In panel C, the arrowheads indicate Sox2+ cells occupying the mantle zone and the asterisk a group of Sox2+ cells abandoning the neural tube. The bar graph in panel D shows the mean ± SD of the Sox2+ and HuC/D+ cells among the transfected population. Lightning bolts denote transfection and no significant statistical differences were observed among the three groups.

